# Development and validation of two economical and flexible immunoassays for detecting antibodies against LCMV in mouse serum

**DOI:** 10.64898/2026.04.07.716918

**Authors:** Rebekah Honce, Jillian German, Ella K. Botten, Chloe Schiff, Emily Van Beek, Allysen Henriksen, Kanayo Ikeh, Akshay Neeli, Philip Eisenhauer, Inessa Manuelyan, Jason W. Botten

## Abstract

Measurement of antibody responses to viral infection is essential for surveillance, diagnostics, epidemiological research, and natural history of infection studies. However, current methods to detect virus-specific antibodies are often resource-intensive and impractical for deployment in outbreak settings or in field-based studies. This manuscript presents two economical, high-throughput immunoassays—the cytoblot immunoassay (CBA) and strip immunoblot assay (SIA)—for detecting and quantifying anti-lymphocytic choriomeningitis mammarenavirus (LCMV) antibodies in mouse serum. To validate, we tested serum from acutely or persistently experimentally infected mice. Both assays detected LCMV-specific IgG and IgM antibodies with high sensitivity and specificity across multiple timepoints. By facilitating the study of immune responses in rodent reservoirs, these tools can enhance our understanding of zoonotic viral transmission, provide scalable platforms for outbreak preparedness, and serve as adaptable models for the development of rapid serological assays for other viral pathogens.

## Introduction

One facet of the immune response to viral infection is the production of virus-specific antibodies. These antibodies play key roles in the humoral response to infection, ranging from direct viral neutralization to complement-mediated and various Fc effector functions (1). The evaluation of antibody serostatus can have practical functions in both viral surveillance (2–7) and in laboratory research (8), such as the assessment of viral exposure, infection, or vaccination in human cohorts or experimental animal infection models, and in the monitoring of infection status and transmission in reservoir species. However, when available, most existing methods to quantify virus-specific antibodies are time- and labor-intensive, requiring expensive detection reagents and specialized equipment, or can only be done at centralized regional or national health laboratories or contract research organizations. This stagnates research progress and hinders local responsiveness during outbreak scenarios, especially for zoonotic viruses.

Viral spillover from animal reservoirs to human or other incidental hosts is a major global health concern highlighted by the predominance of animal-origin viruses on the World Health Organization’s list of priority diseases and the National Institutes of Health list of biodefense pathogens. Mammarenaviruses, ambisense RNA viruses naturally harbored primarily by rodents worldwide, remain a persistent threat to global health as evidenced by recent Lassa virus outbreaks in West Africa (9), periodic spillover of Junín, Machupo, and Guanarito viruses—among others—in South American human populations (10), and emergence of novel pathogenic mammarenaviruses (11). The natural history of viral infection in rodent reservoirs is incompletely understood, especially regarding the immune response to persistent arenavirus infection. For reservoir rodents, understanding historical infections can aid in studies of zoonotic virus prevalence and enable identification of putative hosts. Similar antibody detection approaches, such as those employed in the settings of orthohantavirus or coronavirus infection, have proven effective to ascertain infection history, particularly when paired with molecular epidemiology during viral surveillance of rodents worldwide (12–15).

Here, we report and compare two in-house immunoassays for rapid detection of anti-lymphocytic choriomeningitis mammarenavirus (LCMV) antibodies in the serum of infected experimentally infected mice. Both approaches are economical and high throughput (13, 16, 17). Using the cytoblot immunoassay (CBA), we can qualitatively determine seropositivity of up to 40 samples or quantitate the endpoint titer of up to 10 samples per 96-well plate. The strip immunoblot assay (SIA) test provides a parallel reduction in equipment and supplies needed ideal for field applications. Both assays were tested for the presence and, in the case of the CBA, endpoint titer of LCMV-specific IgG and IgM antibodies in two cohorts of experimentally infected mice. This approach provides a framework for the rapid development of sensitive and specific immunoassays in outbreak scenarios and can be further optimized for field-based applications.

## Results

### Qualitative detection of LCMV seropositivity using an in-house strip immunoblot assay

We first assessed the qualitative presence of anti-LCMV specific antibodies using a strip immunoblot assay (SIA) (13, 16). We tested three antigen sources. First, infected cellular lysates (BHK cells at 48 hours post-infection (hpi) with LCMV Armstrong 53b) were prepared for immunoblotting and loaded onto a 2D-well gel for protein separation. After transfer onto a nitrocellulose membrane, strips approximately 3 mm wide were cut and stored in blocking buffer prior to serum sample addition (**Fig 1A, Supplemental Fig 1A**). Second, cell-free LCMV virions derived from the supernatants of Vero E6 cells 48 hpi with LCMV Armstrong 53b were sucrose-banded, then prepared for immunoblotting prior to electrophoresis (**Supplemental Fig 1B**).

**Fig 1.**
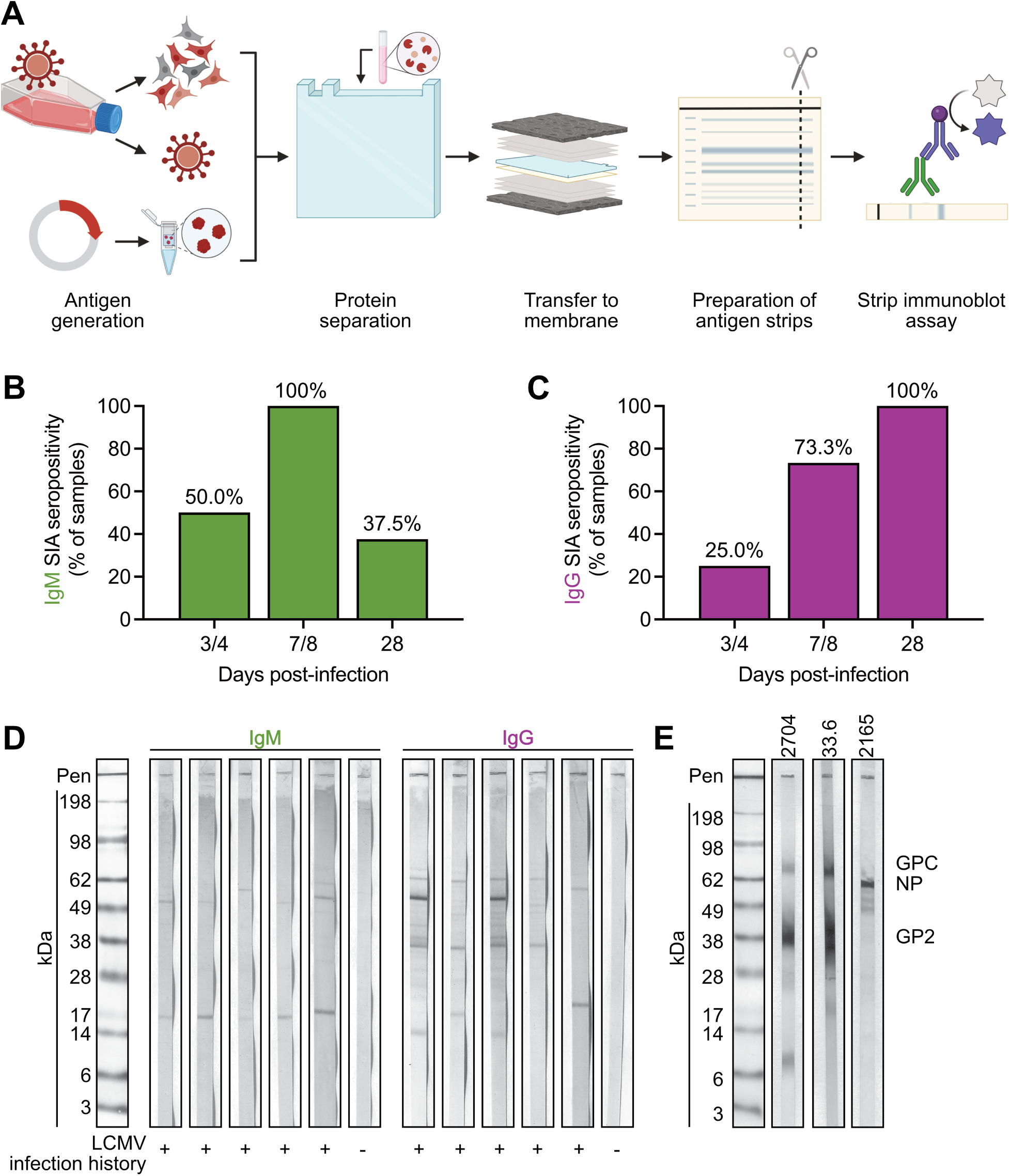
Assessment of LCMV seropositivity by strip immunoblot assay. (A) Preparation of strips containing LCMV antigen derived from BHK whole cell lysates at 48 hpi via gel electrophoresis. (B-C) Adult C57BL6 mice (n=8-20 per timepoint) were intraperitoneally inoculated with 1000 PFUs of Armstrong 53b and blood collected at indicated timepoints. Percentage of samples with (B) IgM or (C) IgG positive tests represented. (D) Representative staining patterns of mice in (B, C). (E) Representative staining of known monoclonal (33.6, αGPC/GP2) and polyclonal antibodies (2704, αGPC/GP2; 2165, αNP) raised against LCMV. Data in (B-C) graphed as percent of total samples.

Finally, purified recombinant LCMV glycoprotein subunit 2 (GP2 aa 266-498) was commercially obtained and prepared as above (**Supplemental Fig 1C**). For assessment of seropositivity, 5 µL of mouse serum from experimentally infected animals was added to each antigen loaded strip in 500 µL of blocking buffer for a dilution of 1:100 and allowed to incubate overnight with rocking at 4°C. After primary sample addition, alkaline phosphatase-conjugated secondary antibodies were added for 1 h at RT. Strips were then washed, and alkaline phosphatase signal was developed. In our initial studies we saw more consistent and strong banding from the whole cell lysate antigen source than either purified viral protein or virions (**Supplemental Fig 1A-C**). Purified virions also showed strong signal, especially in comparison to rGP2; however, due to the labor, cost, and biosafety considerations associated with sucrose purification of high-titer, virus-containing cell culture supernatants via ultracentrifugation, we decided to optimize our assay conditions against whole cell lysates. In attempts to shorten the sample incubation period of the SIA from overnight, we failed to consistently detect distinct banding patterns after a 1-hour incubation with known low titer, positive serum samples while high titer, positive samples were clearly identified regardless of incubation time (**Supplemental Fig 1D**). Regardless of primary antibody incubation time, longer incubations universally and significantly increased band staining intensity (**Supplemental Fig 1E**). Finally, we tested our optimized assay featuring LCMV-infected BHK whole cell lysates against a larger panel of serum samples from experimentally infected adult mice. We consistently detected the presence of both IgM and IgG viral specific antibodies across multiple timepoints following infection, in line with the expected response to acute LCMV infection of an immunocompetent C57Bl/6 mouse with LCMV strain Armstrong 53b (**Fig 1B,C**; representative images in **Fig 1D**). The pattern of antigens recognized varied from mouse-to-mouse, in line with the recognition of multiple LCMV antigens by the polyclonal sera (**Fig 1E**). When stored in the dark, strips remain readable for at least 3 months post-staining with no quantifiable reductions in staining intensity (**Supplemental Fig 1F,G**).

### Endpoint titration of LCMV-specific antibodies in mouse sera using an in-house cytoblot assay

While the SIA could robustly and reproducibly qualitatively detect seropositivity, it proved cumbersome in quantifying endpoint titers due to the large number of strips and reagents required for a single sample. To overcome this limitation, we prepared a cell-based enzyme linked immunosorbent assay (ELISA), or cytoblot immunoassay (CBA), that could qualitatively detect seropositivity but also determine endpoint antibody titer in a more high-throughput manner (17, 18). Vero E6 cells were seeded at a density of 10,000 cells per well of an opaque, white-bottomed 96-well plate and overlaid with viral inoculum containing LCMV at a concentration of 100 focus-forming units per well. The first column of the plate was overlaid with media containing no virus as a negative antigen control (**Fig 2A**). Plates were fixed at 24 hours post-infection and prepared for immunostaining. Pre-diluted (1:100) serum samples from experimentally infected adult mice were added in 1:2 serial dilutions across the plate, and signal visualized after a 1-hour incubation by the addition of horseradish peroxidase-conjugated secondary antibodies and appropriate developing reagent. We were successful in quantifying endpoint IgG and IgM titers across multiple timepoints post-infection (**Fig 2B, C**, representative images **Fig 2D**). Like the SIA, we had no cross-reactivity to serum from control animals. We saw minimal effects of blocking reagent in the CBA (**Supplemental Fig 2A-C**), as there was no significant difference in background staining regardless of blocking reagent used (**Supplemental Fig 2B,C**). Overall, longer sample incubation time increased the strength of antigen staining in the assay while overnight blocking modestly but significantly reduced background staining; however, the shortest assay tested (1-hour block plus 1-hour primary sample incubation time) still yielded clear results in assessing qualitative seropositivity and endpoint titer for the CBA (**Supplemental Fig 2D, E**). Note, as has been previously reported for similar viral focus assays or ELISpot assays, this signal does fade over time which can be slowed via storage in a dark, airtight container at 4°C (data not shown).

**Fig 2.**
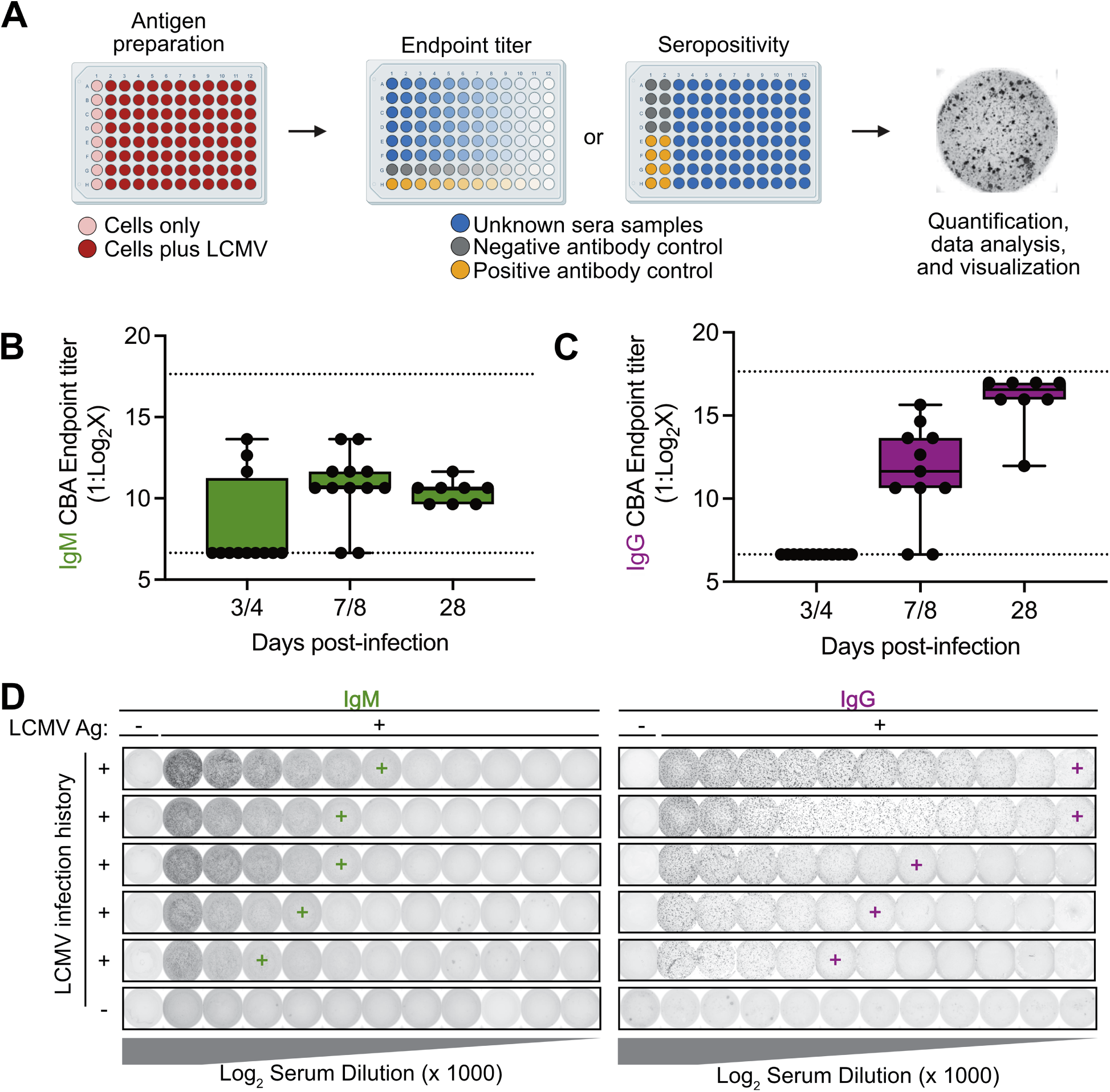
Assessment of LCMV endpoint titer by cytoblot immunoassay. (A) Preparation of test plate using susceptible Vero E6 cells and inoculation with LCMV. (B-C) Adult C57BL6 mice (n=12 per timepoint) were intraperitoneally inoculated with 1000 PFUs Armstrong 53b and blood collected at indicated timepoints. Endpoint (B) IgM or (C) IgG titers of individual mice at each timepoint. (D) Representative staining of samples in (B, C). Data in (B-C) graphed as box plots with whiskers indicating the minimum to maximum values and the center line indicating median values with points representing each sample. Endpoint titers in panels (B-C) were called by eye from trained users. Schematic in (A) designed with bioRender.

### Quantification of endpoint titer using commercially available software

To standardize the assessment of endpoint titer from the CBA, three commercially available software systems were compared to values manually obtained by human eye. Processed plates were scanned, generating the image provided to human scorers and used in method #1 as well as the raw image file required for methods #2 and #3 (**Supplemental Fig 3A-G**). Methods #1 and #2 performed similarly to measures by trained scorers (**Supplemental Fig 3A-D, F**), while method #3 typically under-reported the endpoint titer (**Supplemental Fig 3B, E, G**). Next, we compiled a set of n=30 samples that we have shown to be positive previously in both the SIA and CBA and n=10 known negative samples and provided these materials to trained scorers to mimic how new users may interpret the expected results of this assay (**Supplemental File 1**). All trained users reliably determined seropositivity; however, differences between users resulted in endpoint titers spanning 2 to 6 dilutions for individual samples (**Supplemental Fig 4A-E**). Notably, this variability reflected consistent user-specific calling patterns rather than random error, and the overall standard deviation across positive samples remained low (1.42 dilutions, **Supplemental Fig 4E**). Similarly, test subjects were able to qualitatively assign seropositivity from the SIA with high accuracy when compared to known infectious history (**Supplemental Fig 4F, G**). Accuracy of true positives ranged from 86.7 to 100% (**Supplemental Fig 4H**), with 2 false positives called by the 10 scorers across the 10 known negative samples (**Supplemental Fig 4I**).

### Assessment of LCMV seropositivity in experimental infection models

Next, we tested the performance of both assays against serum samples collected under differing experimental infections (**Fig 3A**). As displayed previously, mice infected as immunocompetent adults had readily detectable IgM and IgG antibodies post-infection in both the SIA (**Fig 1**) and CBA (**Fig 2**). As anticipated, IgM antibodies were detected via SIA as early as day 3 post-infection but waned by day 28 post-infection. In contrast, IgG antibodies were first detected on day 3 post-infection but continued to increase in both frequency of seropositivity and endpoint titer through day 28 post-infection. The detection of antiviral IgG antibodies via CBA in samples collected on day 8 post-infection that were negative by SIA suggests a higher level of sensitivity in the CBA compared to the SIA. We could successfully detect IgG levels in adult mice up to day 75 post-infection (the latest timepoint tested, **Fig 3B**).

**Fig 3.**
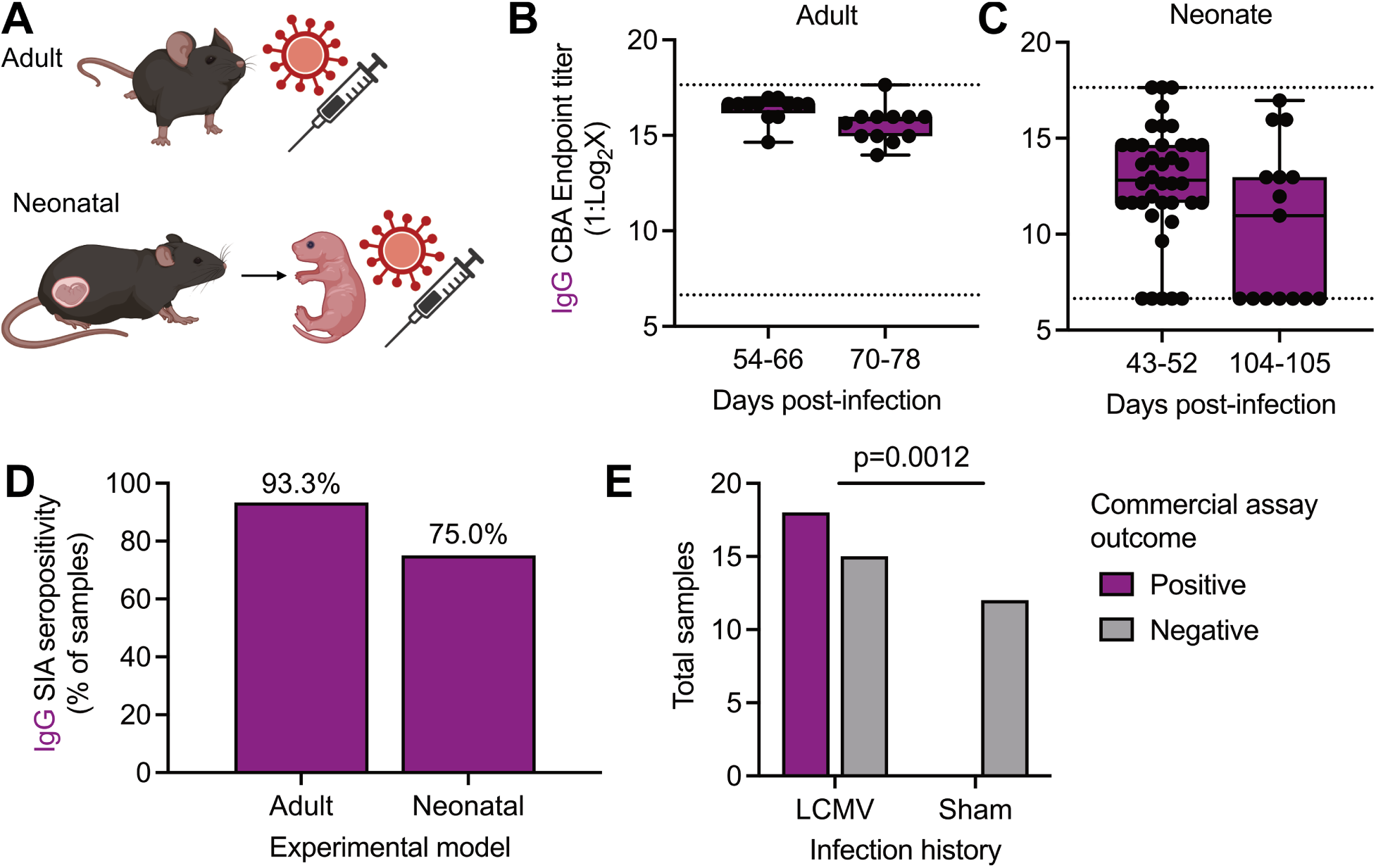
Performance of assays in experimental infection models and against commercially available kits. (A) Experimental schematic of directly infected adult or neonatal, immunocompetent mice. Using the CBA, the (B) endpoint IgG titer of directly infected adult mice at indicated timepoints (n=12 mice per timepoint) and (C) endpoint IgG titer of directly infected neonatal mice at indicated timepoints (n=40 or n=15 mice at d 43-52 pi or d 104-105 pi, respectively). (D) percent SIA-positive sera samples from directly infected adult (n=8) or neonatal samples (n=15) previously found to be positive in B, C). (E) Comparison of n=33 known positive and n=12 known negative samples in a commercially available assay. Data in (B, C) represented as min-to-max boxplots with all points shown, in (D) as percent of total samples, and in (E) as absolute values. Dotted lines indicate the lower limit of detection. Statistical comparison in (B-C) via Welch’s t-test, and in (E) via Fisher’s exact test. Schematic in (A) designed with bioRender.

Next, we compared IgG antibody production in differing experimental model systems. We elected to only probe IgG levels across these systems due to the temporal nature of IgM positivity. LCMV infection in laboratory mice is highly variable based on viral strain, dose, and age of the mouse (19). Thus, in addition to the adult infection model which leads to an acute, self-limited infection in immunocompetent hosts, we also tested a model of persistent infection. Neonatal mice (age day 0) were inoculated via intraperitoneal infection with LCMV to establish a persistent infection. At multiple timepoints through the first 6 months of life, serum was collected via retroorbital bleed, with seropositivity determined via SIA and endpoint titer via CBA (**Fig 3C, D**). A high proportion of directly infected neonatal mice remained seropositive to at least day 50 of life, although endpoint titer did wane, albeit non-significantly, with age (**Fig 3C**). Finally, using randomly selected samples previously determined to be positive or negative in our developed assays, we assessed a commercially available anti-LCMV ELISA (**Fig 3E**). Diluted (1:100) serum samples were further diluted 1:5 for 3 total dilutions per sample. Out of 33 experimentally positive samples (known history of infection and previously positive on either the CBA or SIA), 18 were above the assay threshold, giving a positive predictive value of 0.5455 (95% confidence interval, 0.3799 to 0.7016). We had no false positives in 12 known negative samples tested (**Fig 3E**).

## Discussion

Here, we have described two assays that performed well under experimental conditions to detect seropositivity and determine endpoint anti-LCMV antibody titer in experimentally infected mice. In validation of the CBA and SIA against known experimentally infected adult mice, both assays showed high sensitivity and specificity in detecting antibodies after infection of adult mice. Similar approaches to quantify LCMV- and arenavirus-specific antibody titers have been reported in humans (3, 18) and rodents (4). Past antibody detection assays have reported high levels of specificity and sensitivity from experimentally and naturally infected mice (20), although neonatally infected mice did show reduced intensity of reported positive signals overtime (21, 22). Thus, we limited calculations of sensitivity and specificity to the adult infection model. Here, across a total of n=55 samples assayed in the CBA ranging from day 8 to 77 post-infection, we achieved a specificity of 100% and sensitivity of 95.35%. For n=52 samples assayed in the SIA, we achieved a specificity of 100% and sensitivity of 69.23%. When sensitivity was calculated, inclusion of samples obtained near the cusp of expected seroconversion or extending into the potential timeframe for antibody waning (prior to day 10 and after day 70 post-infection, n=2/20 for CBA and n=5/26 for SIA) may have led to conservative estimates of assay sensitivity (23, 24). Antibody responses to LCMV typically develop after the peak of the early T cell response suggesting samples obtained during this early 3-7 day period following infection may fall below the limit of detection despite ongoing immune priming (25, 26). Observational surveillance studies also show higher positivity frequencies by RT-PCR when assessing viral infection status versus antibody-based ELISAs, suggesting viral carriage and infection status in LCMV-infected *Mus* may not be easily gleaned from assessment of seropositivity alone (27–29). However, our assays outperform a commercial LCMV-specific ELISA commonly used in surveillance studies, even when starting with the same dilution of serum (1:100).

Throughout our studies, we consistently noted the CBA had higher sensitivity than the SIA, potentially due to the greater amount of concentrated LCMV antigens compared to the separated protein(s) used as an antigen source in the SIA. Further, viral foci in the CBA wells yielded a consistent staining pattern across different serum samples, whereas antigen bands in the SIA proved more variable between samples which reduces the ease of interpretation of the SIA. Additional studies potentially should be performed to further optimize the assays for field use, including studies on cross-reactivity with other mammarenaviruses in areas where multiple circulate naturally (13). Although endpoint titers for individual samples spanned a range of 2 to 6 dilutions when determined by different users, this variability was driven largely by consistent, user-specific calling patterns rather than random inconsistency (**Supplemental Fig 4B**). Importantly, the overall standard deviation across positive samples was low, indicating the assay yields reproducible results despite inter-user variation. These findings suggest the assay is inherently robust, and remaining variability likely reflects subjective interpretation of endpoint criteria which may be further minimized through standardized training or automated endpoint determination (**Supplemental Fig 3**).

Beyond analytical performance, practical considerations of these assays are also essential for implementation in the laboratory. In outbreak scenarios, it may be prudent to test for human and animal exposure against the “whole” virus proteome to capture the full range of antigen and antibody pairs instead of relying on a single purified protein. This may be especially crucial in public health emergences when the highly antigenic and immunodominant viral proteins are unknown and rapid detection is more important than protein specificity. These features may also be relevant for detecting LCMV seropositivity in wild-caught and laboratory rodents used for experimental research. For example, screening wild rodents prior to establishing new laboratory colonies can reduce the risk to research staff and may be necessary for specific-pathogen free research goals (6, 7). Further, the increasing use of “dirty” pet store mice in immunological modeling studies necessitates clear screening of animals to safeguard against inadvertent pathogen introduction, LCMV or others, to animal facilities (30, 31). Routine serosurveillance using whole virus-based assays could help identify silent LCMV outbreaks within animal facilities where infections may otherwise go unnoticed due to subclinical presentation in adult animals, but can have serious impacts to human health and study outcomes (32–34). The ability to rapidly deploy cost-effective assays without the need for specialized equipment enhances their utility for colony management, quarantine decisions, and field surveillance, strengthening both research reproducibility and biosafety oversight.

These assays provide a practical framework that substantially reduces development time and cost while maintaining flexibility across laboratory and field settings. With the right combination of susceptible cells and isolated virus in-hand, these assays can be produced in-house from antigen generation to seropositivity assessment with an approximate cost of less than $1.50 USD per sample for CBA and less than $6.15 USD per sample for the SIA (**Table 1**), markedly lower than the per sample cost for recombinant protein production or commercial kits (range $9.37–$14.58). However, the use of recombinant protein provides an advantageous option for facilities with limited cell culture abilities, for lower biocontainment needs, and to limit the cross-reactivity of novel viruses to existing, closely-related strains and thus the flexibility of these methods is a highlight for their application in these settings. Both assays also minimize resource consumption by detecting viral antigens directly on permeabilized monolayers (CBA) or nitrocellulose membranes (SIA), avoiding the need for soluble colorimetric reagents prone to high background staining with polyclonal sera. While soluble reagents enable clear quantification of data, an advantage of the CBA or SIA over a traditional ELISA is that trained visual assessment can reliably distinguish true viral foci or antigen-positive banding from background, particularly when experimental samples are processed alongside known positive and negative controls to mitigate batch variability and when viral stocks are carefully standardized (**Fig 2D** versus **Supplemental Fig 2B, D**). Importantly, the HRP- or AP-based “dry” detection systems eliminate reliance on expensive plate readers and proprietary kits, enabling low-cost, visually interpretable endpoint titers without time-sensitive readouts. The SIA further enhances field applicability due to its portability and minimal cold-chain requirements, though its longer time-to-signal and increased serum input compared to the CBA may limit rapid screening and reduce sensitivity for low-titer samples under shortened incubation times (13, 16). Together, these results demonstrate both assays offer reliable, adaptable, and cost–effective tools for detecting LCMV seropositivity, providing a practical framework for rapid implementation across both laboratory and field surveillance contexts.

**Table 1.**
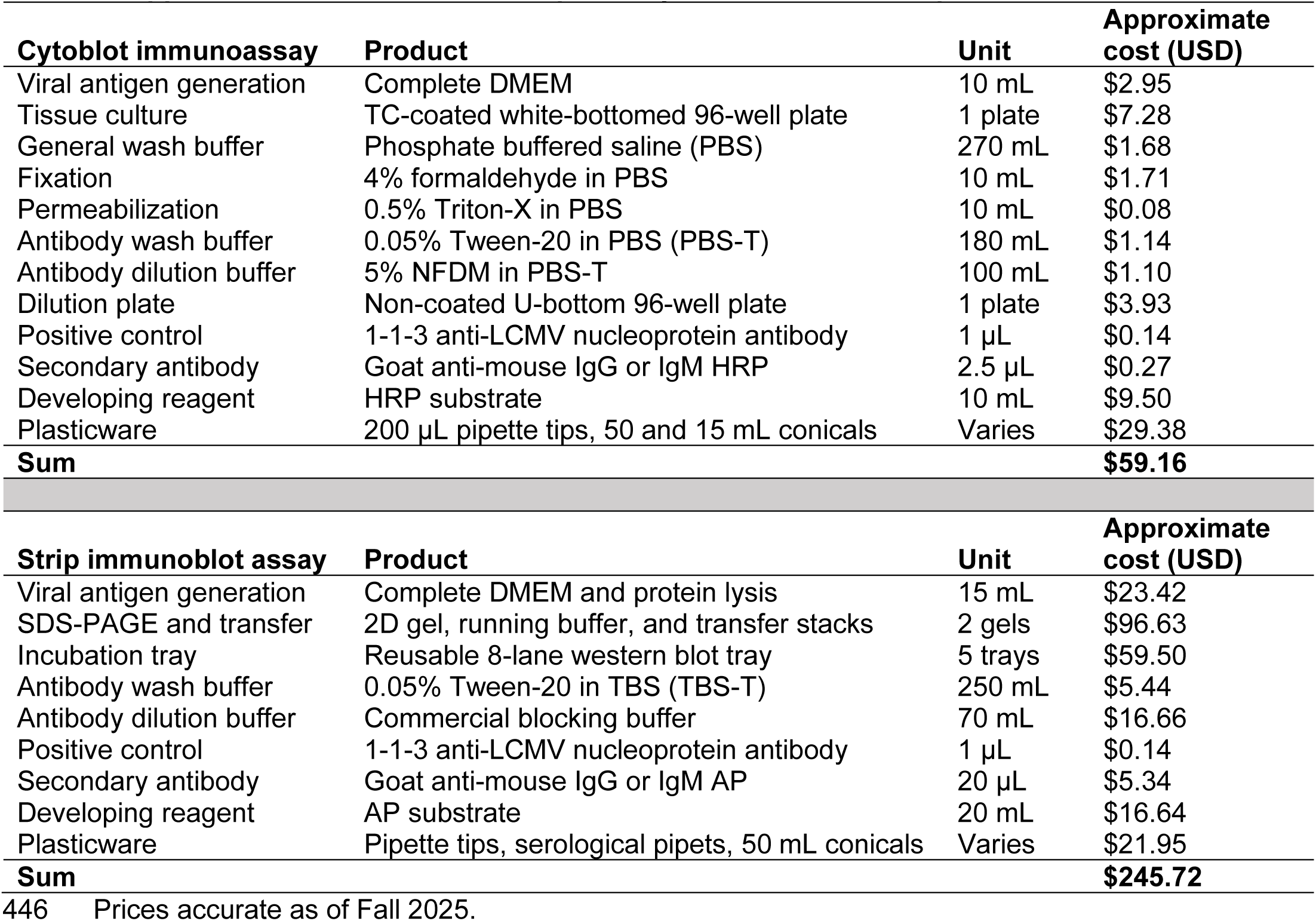
Approximate cost to assess seropositivity in 40 unknown samples.

## Materials and Methods

### Animal husbandry and ethics

Mice were maintained at 12 h light-dark cycles and ambient temperature (27°C) and humidity with food and water provided *ad libitum* for the duration of the experiment. All experiments were conducted in an animal biosafety level 3 laboratory. Investigators were required to wear appropriate respiratory equipment (Versaflo PAPR, 3M). Mice were housed in HEPA-filtered, negative pressure, vented isolation containers and handled only within class 2A biosafety cabinets. All procedures (X2-022) were approved by the University of Vermont Animal Care and Use Committee (IACUC) and complied with the Guide for the Care and Use of Laboratory Animals.

### Generation of serum samples from experimentally infected laboratory mice

Acutely or persistently infected mice were generated as previously reported (19). Briefly, C57BL6 mice were intraperitoneally inoculated with 10 to 10,000 PFUs of LCMV Armstrong 53b diluted in phosphate buffered saline (PBS). To generate persistently infected mice, male and female neonatal mice were inoculated within 24 hours of birth. To induce acute infections that would be cleared within 8-10 days, adult, immunocompetent male and female mice were inoculated between 8 and 12 weeks of age. At indicated timepoints following inoculation, blood was obtained through the retroorbital sinus or terminally via cardiac puncture, allowed to clot at room temperature for 30 minutes, then centrifuged at 2,000 x g for 15 minutes at 4°C to isolate serum. Serum was collected and stored at -80°C until downstream testing. All sera were heat inactivated at 57°C for 20 minutes prior to work on the benchtop.

### Antigen preparation for immunoblotting

All steps were performed in a certified Class II Biosafety Cabinet and with appropriate BSL-2 techniques and personal protective equipment. For whole cell lysates from LCMV-infected cells, baby hamster kidney (BHK) cells maintained in growth medium [DMEM (Gibco #11965084), 10% fetal bovine serum (Gibco #16410-071), 2 mM GlutaMAX (Gibco #35050061), 100 U/mL penicillin and streptomycin (Gibco #15140122)] in T-175 flasks (Corning #431080) were inoculated at MOI=0.01 with recombinant LCMV Armstrong 53b. Plasmids for recombinant virus were generously provided by Dr. Juan Carlos de la Torre (Scripps Research Institute). At 48 hpi, media was removed, cells washed twice with cold PBS, and lysed (5 mM TrisHCl pH 7.4, 150 mM NaCl, 1% TritonX-100, 1 mM EDTA, 5% glycerol, pH 7.5). Lysates were incubated on ice for 20 minutes and clarified through centrifugation at max speed (13,000 rpm) for 10 minutes. Whole cell lysates were stored at -20°C until use. To generate purified LCMV virions, Vero E6 cells were infected with LCMV Armstrong 53b and 2 days later supernatants were collected and subjected to sucrose banding to purify cell-free virions as previously described (35). Recombinant glycoprotein subunit 2 (GP2; amino acids 266-498 from strain Armstrong 53b) was obtained commercially (Creative BioLabs #Vang-0001Lsx) and 20 µg was loaded per 2D-well gel.

### Strip immunoblot assay preparation

To the antigen sources, a 1:1 addition of 4X SDS-PAGE buffer (0.2 M Tris-HCl, 0.4 M DTT, 277 mM SDS, 6 mM bromophenol blue, 4.3 M glycerol) and 10% BME was added prior to boiling at 100°C for 10 minutes. Once antigen sources were denatured and reduced, a volume of 200 µL was loaded into a 2D-well gel (Invitrogen #NP0326BOX) for electrophoresis. After transfer onto nitrocellulose membranes (Invitrogen #IB23002), strips were cut approximately 3 mm wide and stored in blocking buffer (ThermoScientific #37530) for a minimum of 1 hour until probing. Strips were placed in an incubation tray (Biorad #1703902), covered in 500 µL of blocking buffer and 5 µL of mouse serum added to each and incubated overnight at 4°C with rocking. The next day, strips were washed carefully twice with TBS-T prior to addition of alkaline phosphatase conjugated secondary antibody (goat anti-mouse IgG, Invitrogen #31321 or goat anti-mouse IgM, Invitrogen #31326; 1:1000) for one hour at room temperature. After washing, the strips were developed with the addition of a 5-bromo-4-chloro-3-indolyl-phosphate/nitro blue tetrazolium chloride solution (SigmaAldrich #B1911) for 15 minutes at room temperature. To stop the reaction, the developing solution was replaced with ddH2O . Final blots were scanned using a desktop light scanner. For colorfast testing, strips were taped onto standard printer paper and stored in clear plastic sleeve protectors in the dark at room temperature for up to 3 months with images captured monthly and average grey pixels calculated in ImageJ. Quantification is relative to band staining intensity from day of staining. Positive control staining uses various monoclonal and polyclonal antibodies, all used at a 1:1000 dilution (2704 rabbit anti-LCMV glycoprotein (GPC/GP2); 2712 rabbit anti-LCMV GPC/GP1; 33.6 mouse anti-LCMV GPC/GP2; 2165 rabbit anti-LCMV nucleoprotein). Antibodies 2704, 2712, 2165, and 33.6 provided by Dr. Michael J. Buchmeier (University of California, Irvine).

### Cytoblot immunoassay preparation

All steps were performed in a certified Class II Biosafety Cabinet and with appropriate BSL-2 techniques and personal protective equipment. Vero E6 cells maintained in growth medium were trypsinized, counted, and resuspended in growth medium to a concentration of 6.0 x 10⁵ cells per milliliter of medium. A volume of 50 μL was added to each well of a white-bottomed, 96-well tissue culture treated plate (Corning #3524). Infection of individual wells was done immediately after seeding the cells by the addition of a pre-prepared viral inoculum of LCMV Armstrong 53b diluted to 100 focus-forming units in 50 μL growth media per well. Antigen negative wells remained uninfected and were overlaid with only growth medium to serve as a negative control. At approximately 24 hours post-infection, supernatants were aspirated, and cells washed thrice with PBS prior to fixation with 4% formaldehyde by submerging the entire plate. After fixation, plates were removed from the biosafety cabinet for the remainder of the assay. Fixed cell monolayers were washed in PBS then permeabilized via addition of 0.5% Triton-X in PBS for 15 minutes at room temperature. Post-permeabilization, monolayers were washed thrice with PBS then blocked to reduce background signal (see below for blocking reagent optimization). Unknown serum samples were diluted 1:100 in 5% non-fat dry milk (NFDM) in 0.05% Tween-20 in PBS (PBS-T) or indicated antibody dilution buffer. Following incubation of the primary serum samples (see below for incubation optimization), KPL-conjugated secondary antibody (1:2,000, goat anti-Mouse IgG SeraCare #5220-0341 or goat anti-mouse IgM Invitrogen #62-6820) was added for 1 hour at 37°C followed by colorimetric detection (SeraCare #50-78-02) of viral foci. Extensive washing with at least 200 μL PBS-T 4 times each occurred after addition of samples and secondary antibodies. Positive control staining uses antibody mouse anti-LCMV nucleoprotein 1-1-3 purchased from The Scripps Research Institute Antibody Core Facility.

### Optimization of immunoassays

To determine optimal blocking, commercial and in-house blocking reagents were prepared. The following protein blocking reagents were tested: 1% or 5% non-fat dry milk (NFDM); 2% casein (Sigma-Aldrich #C5890 in 1 L buffer [120 mM NaCl, 32.5 mM NaOH, 30.7 mM sodium azide, 7.3 mM Tris-HCL]); 2% bovine serum albumin (BSA), Blocker Blotto (ThermoScientific #37530) and a commercially available blocking reagent (36). All blocking reagents except the ready-to-use commercial regent and 2% casein were dissolved in PBS-T. The following blocking conditions were tested: overnight (12 hours minimum) at 4°C with rocking; 4 hours at room temperature with rocking; and 1 hour at room temperature with rocking. Downstream antigen detection was accomplished as reported above with 1-hour primary sample and secondary antibody incubation at 37°C each. To optimize sample addition, following preparation of the intracellular assay and blocking overnight at 4°C with rocking with 5% NFDM in PBS-T, samples were diluted as above and added to plates. Incubation of the primary samples varied as follows: 1 hour at 37°C, 4 hours at room temperature, overnight at 4°C with rocking. Following extensive washing with PBS-T, downstream antigen detection was completed as reported above.

### Serum antibody titer quantification in laboratory-infected mice

For unbiased endpoint titer quantification, developed plates were scanned (CTL Immunospot S6 Universal M2) and processed in parallel according to the methods outlined below.

Measurements by eye are reported as the mode endpoint titer from 10 blinded and trained individuals. Data are representative of 5 unique CBA preparations which included samples previously determined to be positive (n=30) and negative (n=10) in both the SIA and CBA. Scorers were not involved in sample preparation or assay completion and received scanned copies of all CBA results. Samples were decoded after all results returned. Actual materials provided to individuals are provided in **Supporting Information File 3**.

### Method #1

Scanned images were uploaded to the BioSpot software and quantified using the SmartCount feature. Subjective measures of spot size, separation and background corrections were input based on the using the first positive antibody wells and the negative antigen wells. Using the total intensity of all foreground objects minus background per well, wells were assigned positive if 10-fold greater than the intensity of negative wells.

### Method #2

The individual image outputs from the CTL imager were uploaded to imageJ as an image sequence and converted to 8bit greyscale. Next, thresholding was applied to each well image, using the first positive antibody wells and the negative antigen wells as guidelines for minimum and maximum values. Thresholding was applied using the default method with a light background. On the newly created image stack, the area and mean grey values were measured for each image. Endpoint titer was calculated as the last row with positive signal at least five times greater than background as calculated from negative antigen wells.

### Method #3

Scanned jpeg images were imported into the Image Studio Lite software (v5.5 available at https://www.licor.com/bio/image-studio-lite/) for analysis. The software automatically inverts the image upon import; for ease of analysis red, green, and blue channels were changed to greyscale black on white. Note, the quantification of color intensity does not change based on image brightness, contrast, or color scheme. A region of interest (ROI) was drawn to encompass a single well and the red, green, and blue intensity quantifications recorded for each. Background intensity levels were subtracted from each reading and then values corrected by subtracting the negative antigen well of each row from each test well in the row. Endpoint titer was calculated as the last row with positive signal and is reported individually and averaged across each color channel.

To assess person-to-person variability in seropositivity detection using the SIA, 8 unique strip immunoblot preparations including samples previously determined to be positive (n=30) and negative (n=10) in both the SIA and CBA, were randomized, scanned using a light scanner (CanoScan) and distributed to 10 blinded and trained individuals. Scorers were not involved in sample preparation or assay completion and received scanned copies of all SIA results. Samples were decoded after all results returned. Percent of true positives scored as positive per individual is reported. Actual materials provided to individuals are provided in **Supporting Information File 1.**

### Commercial assay

Serum samples from experimentally infected mice previously scored as positive on both the SIA and CBA were then tested in a commercially available ELISA. This ELISA was specific for murine IgG antibodies against the LCMV nucleoprotein (NP) and is routinely used during LCMV surveillance of wild *Mus* (29). Serum was diluted according to manufacturer’s recommendations (1:5 initial dilution, then 1:20 into provided dilution buffer) for an initial test dilution of 1:100. We further diluted samples 1:5 for 3 total tested dilutions, which were plated in 100 µL. After incubation with detection antibody and developing with provided TMB substrate, absorbance was measured via a plate reader (BioTek). Total absorbance at 450 nm was corrected to the blank, and samples were scored as positive if they exceeded the threshold of the low sensitivity control provided within the commercial kit. Exact assay used is available upon request.

### Statistical analysis, data management, and visualization

Data were collected and stored in Microsoft Excel with statistical analyses and visualization done using GraphPad Prism (v10). Statistical test specifics can be found in individual figure legends. Calculations of sensitivity, specificity, and positive predictive values were done using MedCalc statistical software. Final figures were prepared in Affinity Designer (Serif, v2). Experimental schematics were designed in bioRender and used under a CC-BY 4.0 license.

**Supplemental Fig 1.**
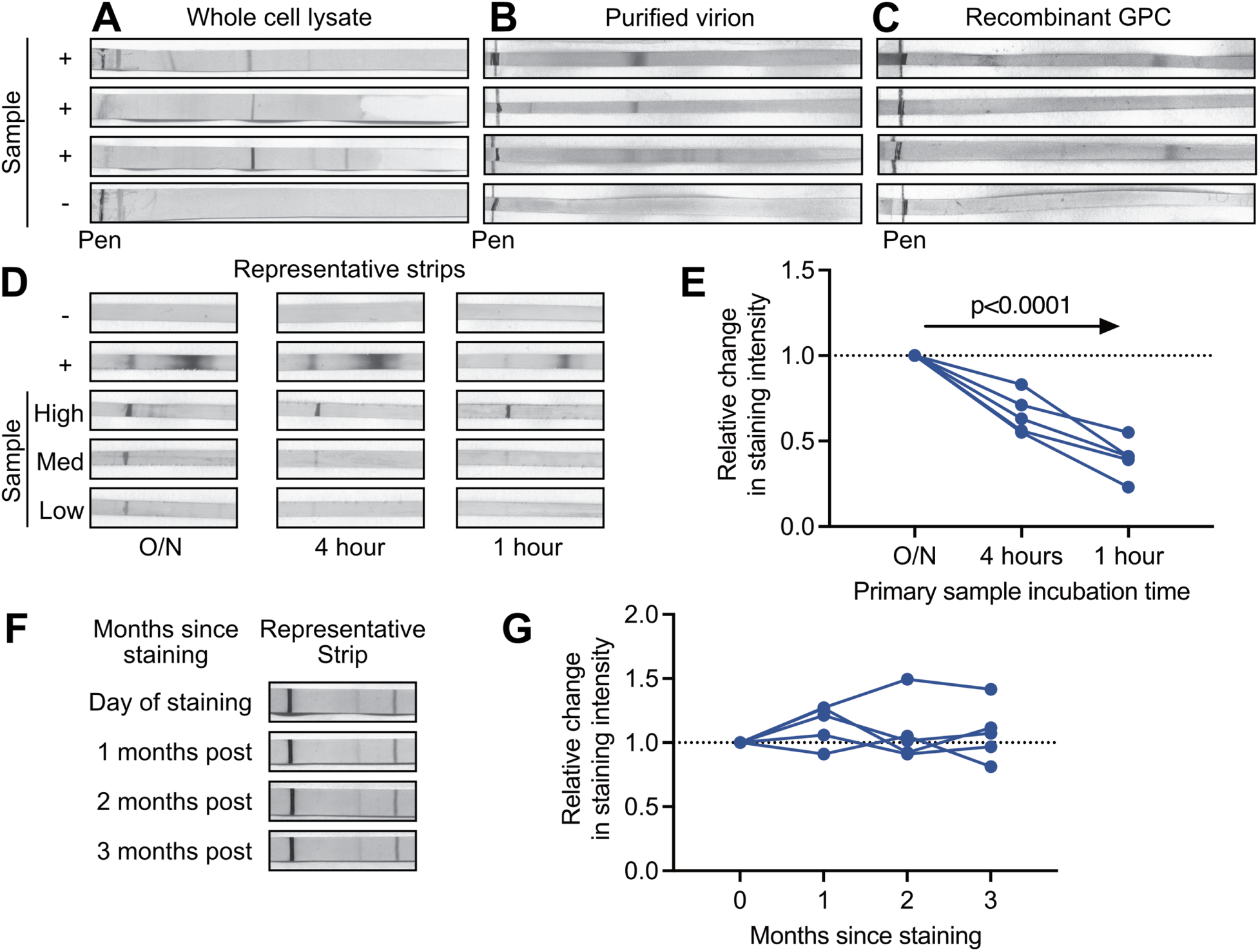
Testing of strip immunoblot conditions. (A-C) Comparison of expected results from western blot membrane strips containing either (A) whole cell lysate from LCMV Armstrong 53b-infected BHK cells at 48 hpi, (B) sucrose banded LCMV Armstrong 53b virions derived from Vero E6 cells, or (C) purified recombinant glycoprotein (GPC) (aa 266 to 498 of strain Armstrong 53b). In each case, strips were exposed to serum from an uninfected, seronegative mouse (-) or from mice infected at 8 weeks of age and sampled at 28 dpi (n=3 independent samples). (D-E) Results of blocking duration optimization SIA testing with (D) showing representative staining for overnight, 4 hours, and 1 hour blocking incubations while (E) quantifies results as connected absolute value data points per sample. Positive control staining was done with 33.6, a known antibody specific for the LCMV glycoprotein (33.6, GPC/GP2) used at 1:500. SIA test in (D) used serum samples with a known high, medium, and low endpoint titer from previous CBA tests. (F-G) Colorfast test of SIA staining with representative strips from day of staining, plus 2, 4, and 6 months-post staining. (F) Quantification of (E) compared to intensity of same band from day of staining using average grey pixels measure in ImageJ. Data represented in (E, G) as connected absolute values for each sample. Statistical comparisons made in (E, G) as a one-way ANOVA test for linear trend. All quantification of staining intensity assessed using the average grey pixels measure in ImageJ.

**Supplemental Fig 2.**
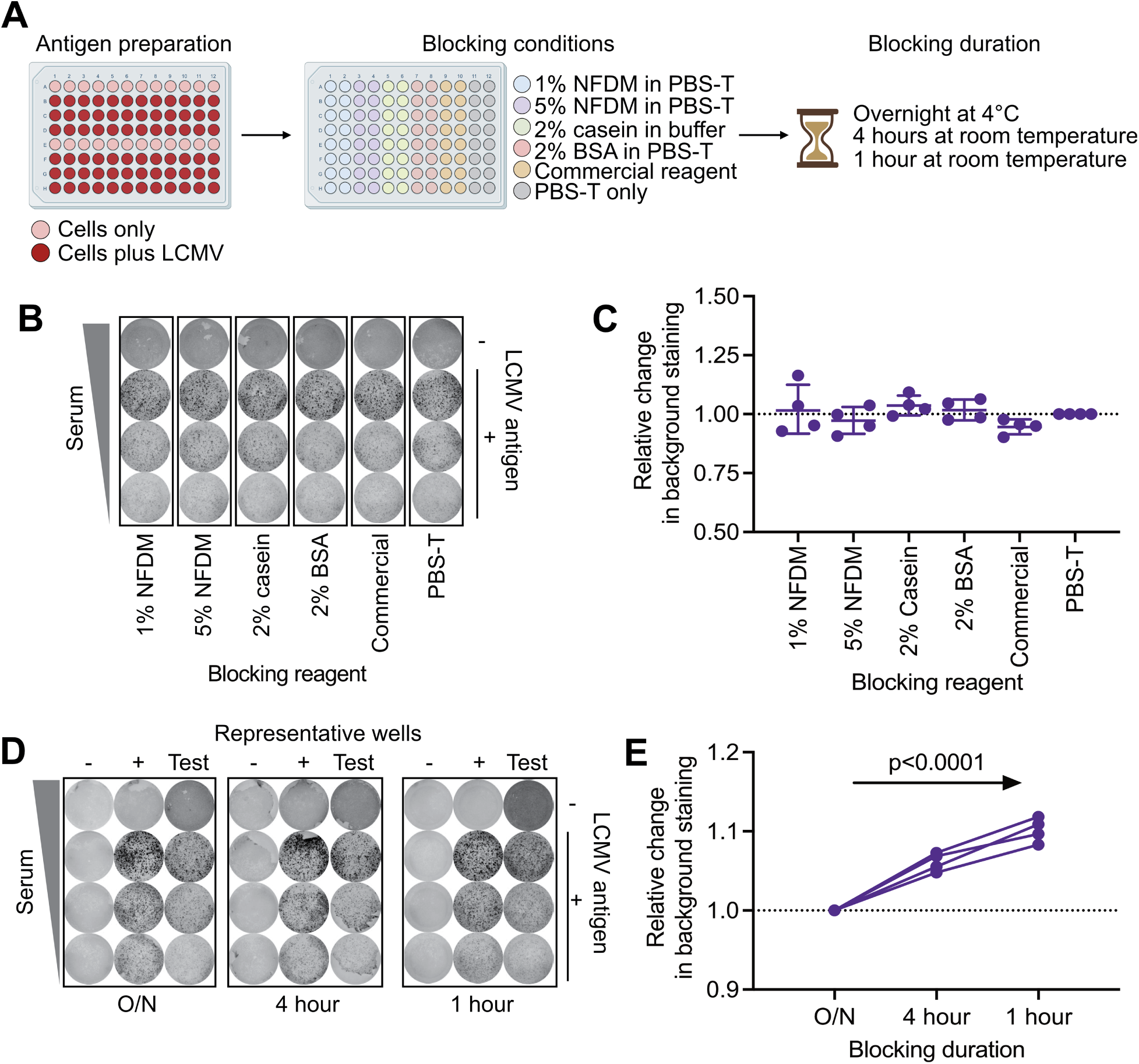
Optimization of cytoblot immunoassay. (A) Test of blocking conditions for CBA. (B) Representative comparison of blocking reagents in CBA using serum from a known positive mouse. (C) Quantification of relative staining antigen negative wells in (B) represented as means ± SEM. (D-E) Results of blocking duration for CBA with (D) showing representative staining for overnight, 4 hour, and 1 hour blocking while (E) quantifies results as connected absolute values data points per sample. Positive control staining is a known LCMV antibody for the nucleoprotein (1-1-3, 1:1000 starting dilution). Negative control staining is phosphate buffered saline. Data represented in (C) as individual datapoints and in (F) as connected absolute values for each sample. Statistical comparisons made in (C) using one-way ANOVA with Dunnett’s multiple comparisons test setting PBS-T as control and in (E) as a one-way ANOVA test for linear trend. All quantification of staining intensity assessed using the average grey pixels measure in ImageJ. Schematic in (A) designed with bioRender.

**Supplemental Fig 3.**
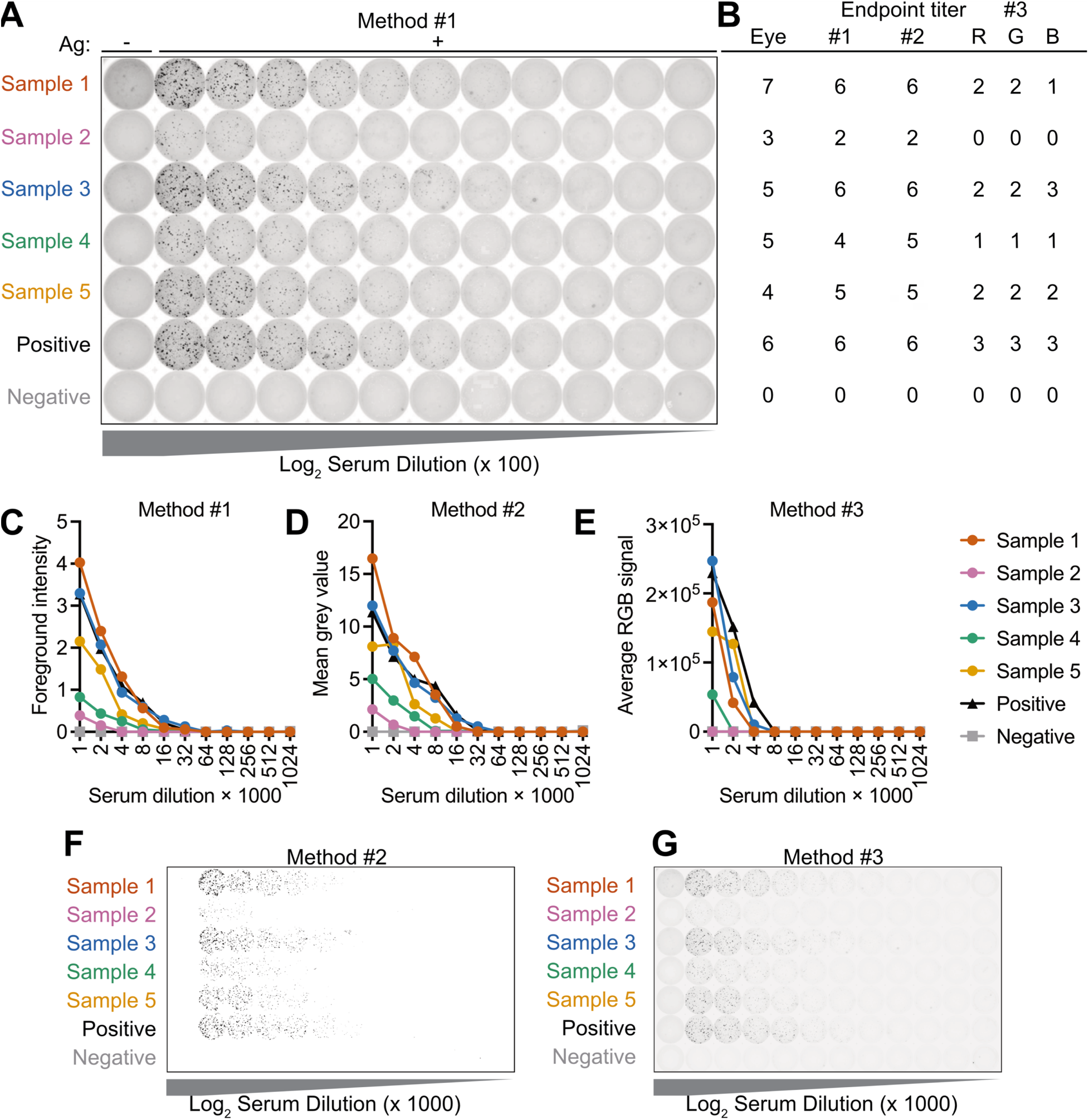
Comparison of seropositivity and endpoint titer calculation methods. (A) Stained CBA plate with representative samples showing a variety of endpoint titers. (B) Comparison of endpoint titer calculation by eye or with automated methods. (C-E) Quantification of staining as represented by images in (A), (F), or (G), respectively. (C) Method #1 calculates the relative intensity of identified foreground objects in each region of interest in (A). (D) Method #2 calculates total black pixels in a region of interest in (F). (E) Method #3 separates the image data in (G) into red, green, and blue channels which are then quantified by the software. Data in (C-E) is graphed as absolute values for each sample. Positive control is staining with a known LCMV antibody (1.1.3, 1:1,000 starting dilution). Negative control staining is phosphate buffered saline.

**Supplemental Fig 4.**
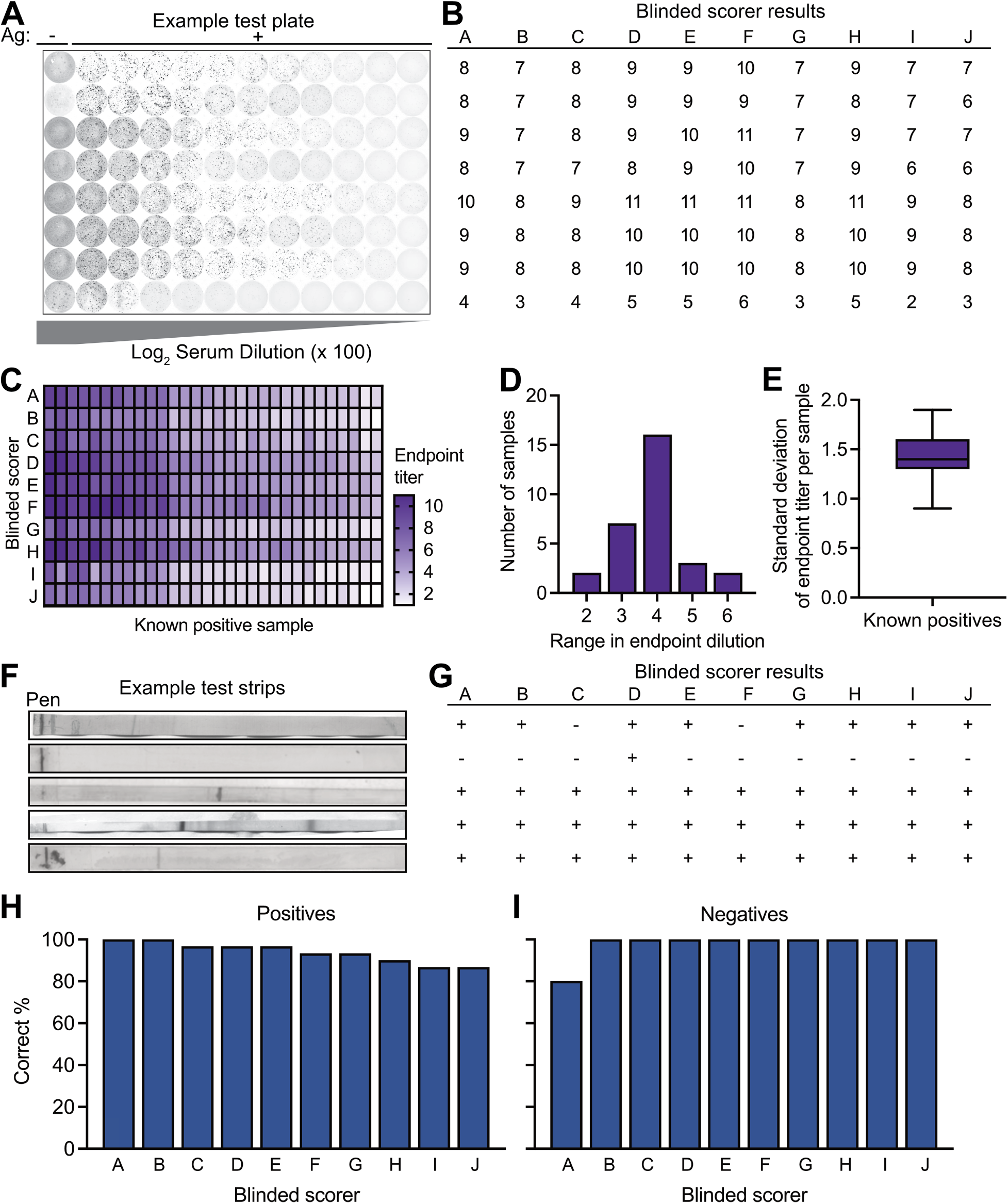
Testing of assay accuracy in trained individuals. (A) Example of test CBA plate provided to trained scorers with (B) individual raw results. (C-E) Variability of endpoint titer determination across 30 known positive samples in the CBA with (D) representing the range in endpoint titer and (E) indicating the standard deviation of each sample endpoint titer. (F) Example of n=5 test SIA stripblots provided to trained scorers with (G) individual raw results. (H, I) Percentage of known (H) positive and (I) negative samples correctly identified by each blinded scorer. Data graphed in (D) as a histogram of range in endpoint titer for each positive sample, in (E) as box plots with whiskers indicating the minimum to maximum values and the center line indicating median values and in (H, I) as percent correct. Trained scorers (n=10) were blinded to sample identification prior to assessment and did not participate in the completion of the assays. Each individual scorer is designated with a letter-labeled column in panels (B), (G), and (H-I).

## Acknowledgements

Author roles are as follows: RH: Conceptualization, Investigation, Methodology, Validation, Formal Analysis, Visualization, Writing - Original Draft Preparation, Writing – Review & Editing. JG: Investigation, Methodology, Visualization, Formal Analysis, Writing – Original Draft Preparation. EKB: Investigation, Methodology, Validation, Formal Analysis. EVB: Investigation, Methodology, Validation. CS: Investigation, Methodology, Validation. AH: Investigation, Methodology, Validation. KI: Investigation, Methodology, Validation. AN: Investigation, Validation. PE: Investigation, Validation. IM: Investigation, Validation, Writing – Review & Editing. JB: Conceptualization, Methodology, Formal Analysis, Writing – Review & Editing, Funding Acquisition, Supervision.

The authors collectively thank Virginia Harget, PhD, Victoria Meliopoulos, PhD, R. Chris Skinner, PhD, Deena Snoke, PhD, and Elena Copson for technical expertise in seropositivity assessment and titer quantification; Jean Celli, PhD and Maria Sonia Godoy-Tundidor, PhD at the University of Vermont and Joyce Oetjen, PhD at the Vermont Department of Health Laboratory for assistance with A/BSL3 facility logistics; Karolyn Lahue, Jennie Bonica, Barbara Jerger, Bailie McDonald, Savannah Godbey, Joshua Woolsey, DVM, and Ida Washington, DVM, PhD at the University of Vermont Office of Animal Care and Management for colony husbandry and health monitoring.

